# The PHB Granule Biogenesis Pathway in *Caulobacter*

**DOI:** 10.1101/2023.07.06.548030

**Authors:** Edward A. de Koning, Mayura Panjalingam, Jessica Tran, Michael R. Eckhart, Peter D. Dahlberg, Lucy Shapiro

**Affiliations:** Department of Developmental Biology, Stanford University School of Medicine, Stanford, California, USA; Stanford Protein and Nucleic Acid Facility, Stanford University School of Medicine, Stanford, California, USA; Division of CryoEM and Bioimaging, SSRL, SLAC National Accelerator Laboratory, Menlo Park, California, USA

**Keywords:** *Caulobacter*, bacterial organelle, PHB granule, PHB polymerase, phasin

## Abstract

PHB granules are bacterial organelles that store excess carbohydrates in the form of water-insoluble polyhydroxybutyrate (PHB). The PHB polymerase, phasin (a small amphipathic protein), and active PHB synthesis are essential for the formation of mature PHB granules in *Caulobacter crescentus*. Granule formation was found to be initiated by the condensation of self-associating PHB polymerase-GFP into foci, closely followed by the recruitment and condensation of phasin-mCherry. Following the active synthesis of PHB and granule maturation, the polymerase dissociates from mature granules and the PHB depolymerase is recruited to the granule. The polymerase directly binds phasin *in vitro* through its intrinsically disordered N-terminal domain. Thus, granule biogenesis is initiated and controlled by the action of a PHB polymerase and an associated helper protein, phasin, that together synthesize the hydrophobic granule content while forming the granule’s protein boundary.

**Importance:** Like eukaryotes, bacteria organize their cytoplasm in subcellular compartments. These bacterial compartments can be membrane-bound (e.g. magnetosomes), or membraneless with protein-encased shells (e.g. carboxysomes). Here we investigate how *Caulobacter* forms membrane-less compartments that store the water-insoluble carbohydrate polymer polyhydroxybutyrate (PHB). A PHB polymerase is essential for granule biogenesis and we observed a direct interaction with the granule associated protein phasin through the disordered N-terminus of the polymerase. We found that PHB granules form by sequential recruitment of key proteins, beginning with the polymerase, and that the granule composition changes as these organelles mature.

## Introduction

The compartmentalization of biochemical pathways in membrane bound subcellular organelles is commonly observed in eukaryotic cells. Although membrane bound compartments are not common in bacteria, there are instances of specific functions attributed to bacterial subcellular compartments such as membrane bound magnetosomes (1) that sequester chemical processes. Other bacterial subcellular organelles that are not membrane bound, include protein encapsulated carboxysomes (2) and those that store polyhydroxy butyrate, PHB (3). These granules are commonly found in many bacterial species, including *Caulob*a*cter crescentus* (4). Consumption of PHB is increased under carbon starvation, suggesting that these granules function as carbohydrate storage organelles (5). Here, we have asked how a new organelle is formed and identified the steps required for the *de novo* formation of *Caulobacter* PHB granules.

Polymers of PHB are hydrophobic and highly insoluble in water. Due to its hydrophobicity, accumulated PHB polymers could function as both the content of the organelle, and as its selective boundary. However, Cryo-EM images have shown that this boundary is composed of a discontinuous layer that is likely proteinaceous (6). Indeed, purified granules have been found to consist mainly of PHB (>95%) and associated proteins (7, 8) and are generally assumed to be membrane/lipid free (9). For an overview of PHB granule composition, see Figure 1A.

**Figure 1.**
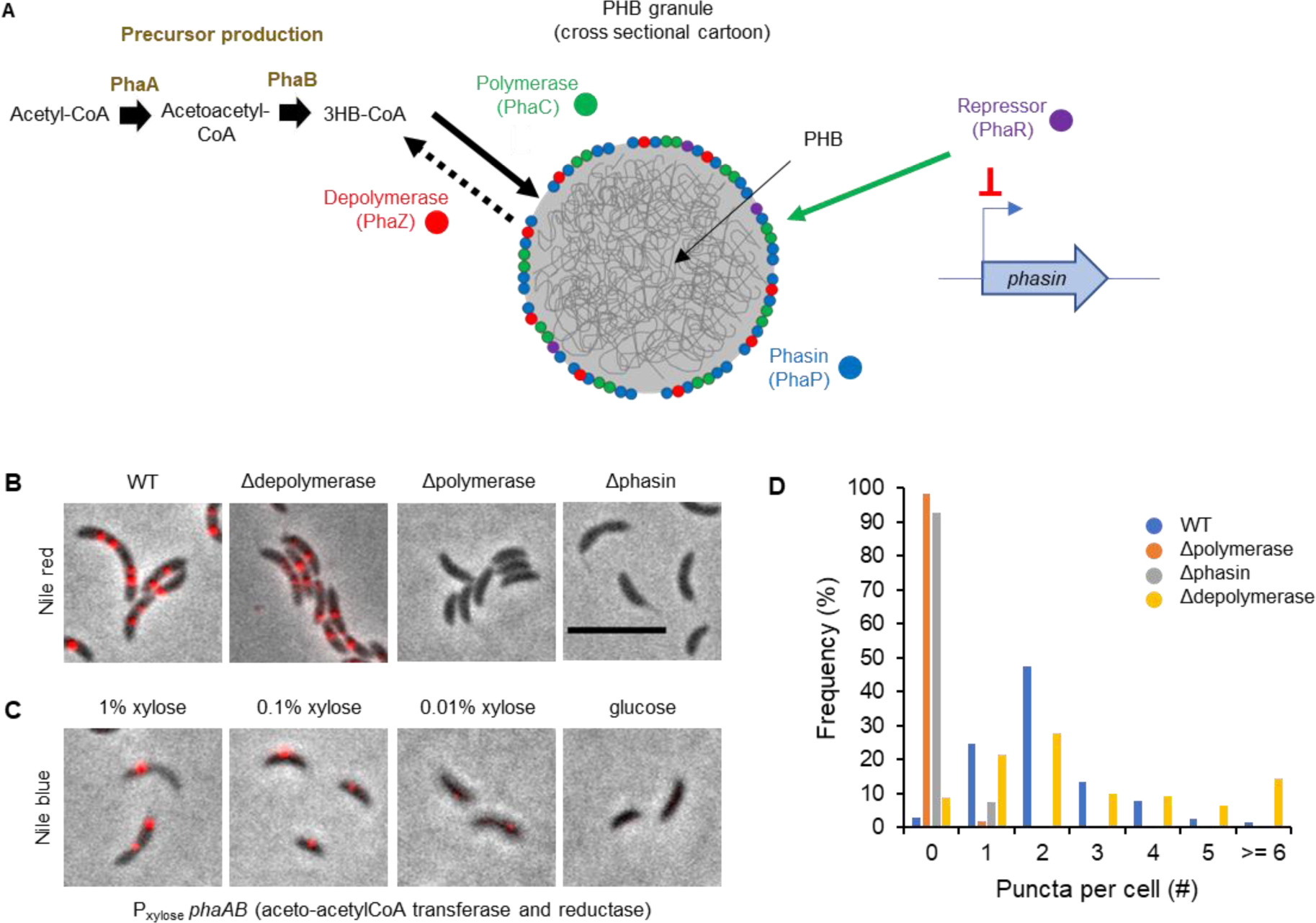
WT PHB granule formation requires both active polymerase and phasin. **A)** chematic of PHB granule organization. A PHB granule consists mainly of water-insoluble PHB, with granule associated proteins on its surface (7). PHB is synthesized by the polymerase PhaC using the 3HB-CoA precursor. This precursor is generated from acetyl-CoA by PhaA and PhaB (23). PHB is consumed by the depolymerase PhaZ, possibly with uncharged CoA as an acceptor for released monomers. Both PhaC and PhaZ are associated with the granule, while PhaA and PhaB are not (8). Another abundant granule associated protein is phasin (PhaP). Its expression is regulated by the transcriptional repressor PhaR, which itself is sequestered to the granule upon PHB synthesis (13, 14). **B)** Deletion of either polymerase PhaC or phasin PhaP results in a loss of nile-red stained granules. Overlay images of phase contrast and Nile-Red fluorescence. Cells are stained in PBS with 1 μM nile red for 20 minutes. Strains used are WT, ΔphaZ (LS5811), ΔphaC (LS5813) and ΔphaP (LS5812). Scale bar is 5 μm. **C)** Reducing precursor availability results in a reduction in nile blue signal and loss of stained granules. Overlay images of phase contrast and nile blue fluorescence. Cells are stained M2G with 10 μM nile blue for 20 minutes. The strain contains a xylose inducible promoter at the start of the CCNA_00544-00545 (phaA-phaB) operon (LS5814). Strain LS5814 was grown for 5 hours in the presence of xylose % as indicated. **D)** Quantification of (B), at least 600 cells were analyzed per strain.

The most abundant granule associated protein is phasin (PhaP), a 15-25 kDa helix-turn-helix protein (10–12). Expression of phasin is regulated by the transcriptional repressor PhaR (13–15). Helical wheel predictions of phasin suggest that the protein is amphipathic, which would position the protein on the surface of the granule. It has been reported that phasin is localized to granule surfaces and can encapsulate purified PHB *in vitro* (10, 16, 17). The crystal structure of *Aeromonas hydrophila* phasin was recently solved revealing assemblies of phasins in stacked hairpins such that the hydrophobic surfaces are shielded from the solution, although it is not known if this conformation occurs *in vivo* on the surface of granules (18). Another protein predicted to reside on the granule surface is the PHB polymerase (PhaC) that catalyzes the polymerization of precursor monomers into PHB (19–21). The precursor monomer is generated in two steps from acetyl-CoA by PhaA and PhaB to make (R)-3-hydroxybutanoyl-CoA (3HB-CoA) (Fig 1A) (19, 22, 23).

PHB is in a crystalline state when synthesized *in vitro*, chemically extracted from PHB containing bacteria, or found in the soil. In contrast, PHB in cells or in gently extracted granules is amorphous (24–27). Bacterial PHB hydrolases can only degrade crystalline PHB (28). Amorphous PHB is specifically degraded by a PHB depolymerase (PhaZ) (29), which is found associated with mature PHB granules (30, 31). The mechanism that keeps PHB in an amorphous state *in vivo* is unknown.

In this study we show that granule formation is dependent on the PHB polymerase (PhaC), phasin (PhaP) and on the active accumulation of the PHB polymer. Granules are formed *de novo* in *Caulobacter* by the initial condensation into foci of the PHB polymerase. The active polymerase catalyzes the accumulation of PHB from the 3HB-CoA precursor. Further accumulation of PHB is dependent on recruitment of phasin to the immature granule, potentially by the direct interaction of polymerase and phasin, which we have shown occurs *in vitro*. Granule biogenesis is stopped by departure of polymerase from the newly formed granule, coincident with the recruitment of PhaZ depolymerase. Taken together, our observations describe how the PHB granule is initiated and changes in composition during its formation and maturation.

## Results

### PHB granule formation requires both an active polymerase and phasin accumulation

PHB synthesis is dependent on the acetyl-CoA acetyltransferase PhaA (CCNA_00544) and acetoacetyl-reductase PhaB (CCNA_00545) to produce the 3HB-CoA precursor from acetyl-CoA (23). These genes have not been previously described in *Caulobacter* but are annotated based on their homology. In *Caulobacter*, the polymerase PhaC (CCNA_01444) and the depolymerase PhaZ (CCNA_00250) have been described previously (4, 32). For an overview of PHB associated proteins, see Fig 1A. The only homologue to phasin found in *Caulobacter* is CCNA_02242, or PhaP from here on. To validate these candidates, we made use of hydrophobic dyes such as Nile Red and Nile Blue that have previously been reported to stain for PHB in a wide variety of bacteria (33–35). We were able to stain PHB granules directly with Nile Blue in the media, and with Nile Red after a PBS wash. We observed fluorescent puncta in *Caulabacter* for both dyes, indicating the presence of PHB granules. These puncta were absent in a mutant lacking the polymerase *phaC* gene and could be rescued by ectopic expression of Polymerase-3xFLAG from a xylose inducible locus (Figs 1B and S1). In cells where the transcription of *phaA* and *phaB* was controlled by a xylose inducible promotor, we could tune the amount of available precursor. By reducing the amount of PhaA and PhaB expression we observed a clear reduction in Nile Blue signal (Fig. 1C and 2A). This indicates that not just the presence, but the activity of the PhaC polymerase is necessary for the Nile Blue puncta. Deleting the phasin PhaP homologue in strain LS5812 also affects puncta formation, as the number of puncta was reduced by 90% in this strain, which was rescued upon ectopic expression of *phaP* from a xylose inducible locus (Figs. 1D and S1). Deletion of the depolymerase did not affect the number of puncta per cell (Fig. 1B).

**Figure 2.**
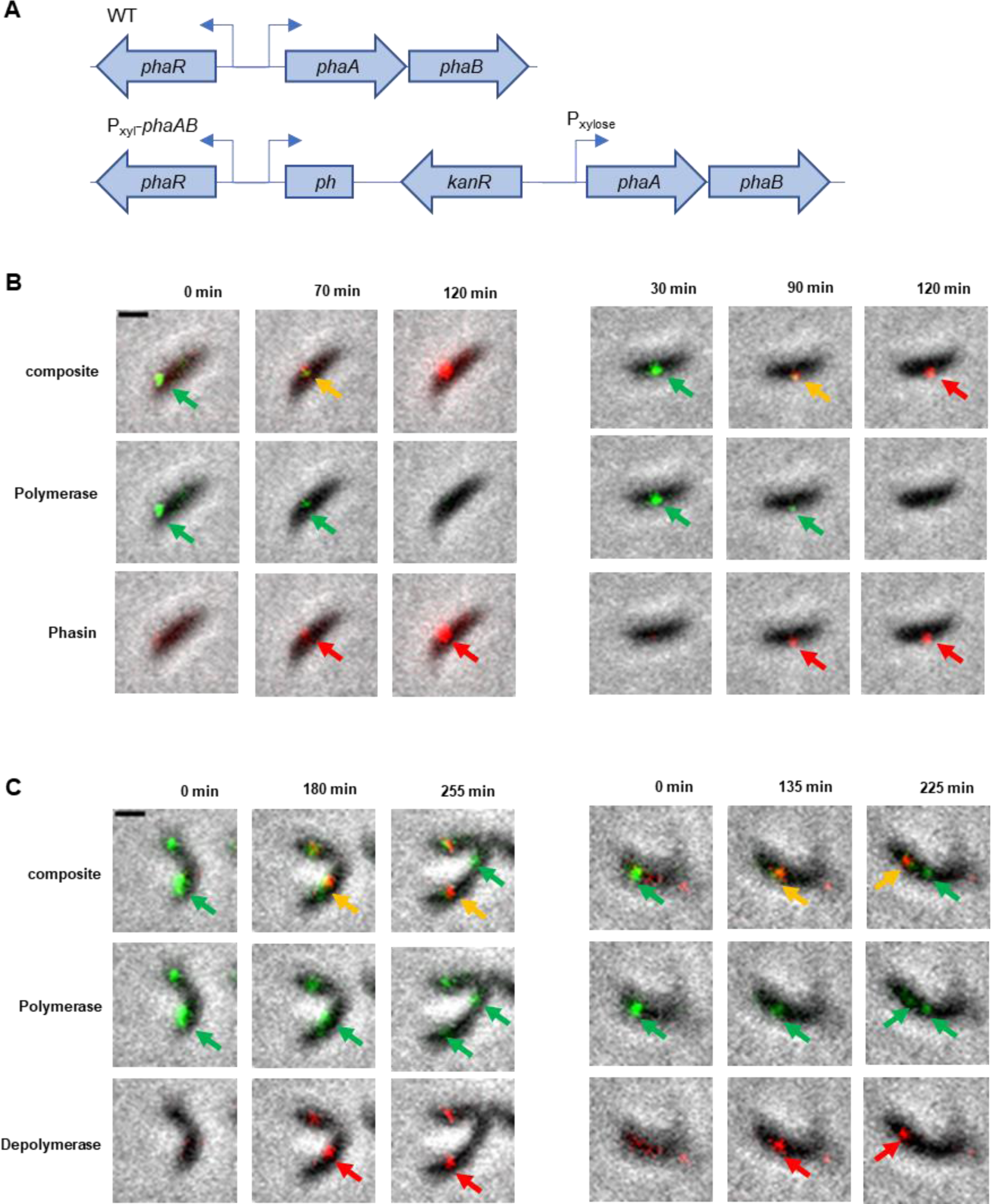
Polymerase condensation precedes phasin and depolymerase condensation. **A)** Representation of the xylose inducible *phaAB* operon (strain LS5814). The genomic region containing *phaR* (CCNA_00543), *phaA* (CCNA_00544) and *phaB* (CCNA_00545) is shown for the WT (top). The xylose inducible promoter was introduced by single crossover integration targeting the 5’ 800bp of *phaA*, leading to a truncated duplication controlled by the WT transcription start site, while the full *phaAB* operon is under control of the xylose promoter. **B)** Timelapse of granule formation. Granule formation was induced by the addition of 1% xylose just prior to timelapse microscopy. Granule formation was visualized by fluorescence imaging of polymerase-GFP and phasin-mCherry (strain LS5871). Condensation of Polymerase is indicated by a green arrow, condensation of phasin by a red arrow, and co-localization by a yellow arrow. Scale bar is 1 μm. **C)** Timelapse of granule formation following glucose introduction to glucose-starved cells. Granule formation was visualized by fluorescence imaging of polymerase-GFP and depolymerase-mCherry (strain LS5821). Condensation of polymerase is indicated by a green arrow, condensation of depolymerase by a red arrow, and co-localization by a yellow arrow. Scale bar is 1 μm.

To further characterize the polymerase and phasin mutants, we imaged *Caulobacter* strains using cryo-electron microscopy (cryo-EM). Analysis of Cryo-EM images of WT *Caulobacter* cells show distributions of both small granules, consisting of poly-phosphate and PHB granules, and large granules, which are exclusively PHB granules (Fig. S2 and Supplementary Text). We observed that the deletion of *phasin* leads to a drastic reduction in larger PHB granules, while simultaneously increasing the number of smaller (< 250nm diameter) granules as compared to WT (Fig. S2). We cannot discern if the observed increase in smaller granules in phasin deletion strains is due to an increase in poly-phosphate granules, or the inability of smaller PHB granules to mature to larger PHB granules. We favor the latter possibility; that we are observing a phasin-dependent maturation of smaller (<250nm diameter) Nile Red negative, to larger (>250nm diameter) Nile Red positive PHB granules.

### Polymerase accumulation initiates granule biogenesis

We used two separate approaches to study time dependent granule biogenesis. i) We controlled the expression of precursor production by placing the phaA and *phaB* genes under the control of the inducible xylose promoter (strain *P_xyl_-phaAB*, Figs. 1C and 2A). In the absence of xylose, granule biogenesis was prevented. While *P_xyl_-phaAB* cells grown without xylose do not have any Nile Blue stained granules, within 1 hour of xylose induction we clearly observe Nile Blue stained granules (Fig. S3A). ii) Cells starved for glucose for at least 24hrs results in a complete lack of Nile red stained granules (Fig. S3B), presumably due to the consumption of all PHB. Reintroducing glucose in the medium restored Nile red-stained granules within 3 hours (Fig. S3B). In each case, controlling precursor production or recovery from starvation, provides a window of time to study *de novo* granule formation in *Caulobacter*.

Using timelapse microscopy, we followed polymerase-GFP and phasin-mCherry signal condensation upon xylose induction of PhaAB expression in a strain containing both fluorescent constructs (Fig. 2B). At the start of the experiment, coincident with the addition of xylose, clear polymerase-GFP condensation is observed, while only faint phasin signal can be detected. After 70 - 90 minutes of PhaAB induction, phasin-mCherry clearly co-localizes with the polymerase. While the phasin-mCherry signal continues to accumulate as the granule matures, the polymerase signal disappears after 2 hours of PhaAB induction.

The departure of the polymerase from a newly formed granule appears to coincide with the mature granule’s final size. We asked if this departure coincided with the recruitment of the depolymerase, which is necessary for granule consumption. In cells recovering from starvation, upon the addition of glucose at time 0, polymerase-GFP puncta were observed, but no depolymerase-mCherry signal (Fig. 2C, t=0). The depolymerase-mCherry condenses at the location of polymerase-GFP localization (Fig. 2B, t=135-180). The polymerase-GFP signal continues to decrease at the first granule as depolymerase-mCherry signal increases, and additional polymerase-GFP signal will condense to initiate the next granule formation (Fig. 2B, t=225-255). These results suggest that PHB granule composition changes over time, initiated with polymerase condensation, followed by phasin accumulation and finally by the recruitment of the depolymerase and loss of polymerase in older granules.

To ascertain that the polymerase condenses into loci prior to PHB accumulation, we used cryo-CLEM (correlative light and electron microscopy) (36–38) with cells expressing GFP-tagged PHB polymerase and stained with Nile Red to image PHB granules (Fig 3A). As shown in Fig 3A, Nile Red exclusively stained larger granules with a diameter over 250nm, and Nile Red intensity positively correlated with granule size (Fig. S4A). We observed that polymerase-GFP localized to both smaller Nile Red negative and larger Nile Red positive granules (Fig. 3 and S4B) and was also observed independent of any granule. Thus, condensation of polymerase precedes the accumulation of PHB polymer as detected by Nile Red. It should be noted that some PHB is likely present in the smaller Nile Red negative granules, which could be below the detection of Nile Red staining. Some membrane deformation can also be observed in these Nile Red stained cells that are absent in unstained cells (see Fig. S2A), possibly due to the PBS wash necessary for consistent Nile Red staining in our experiments.

**Figure 3.**
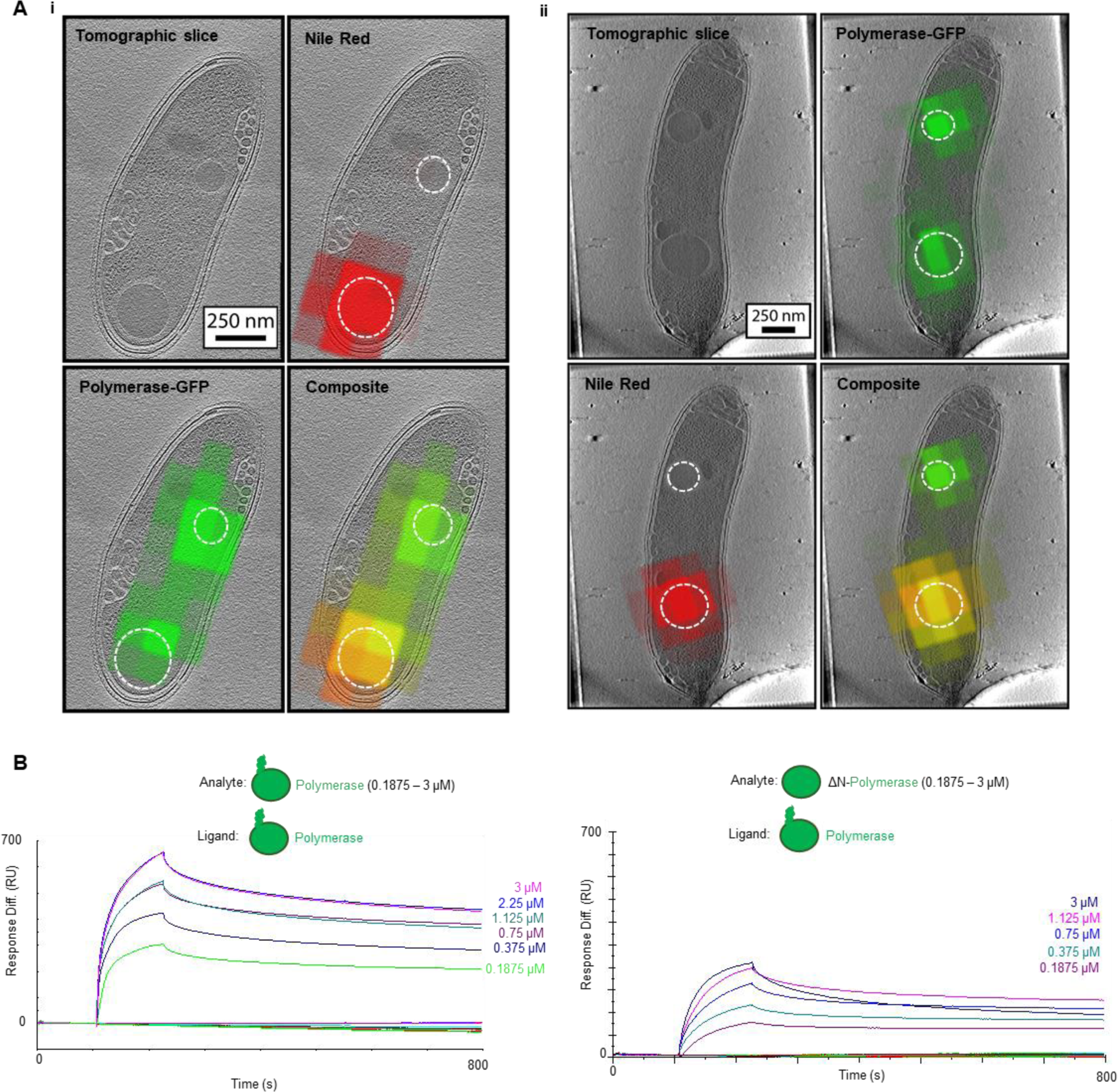
Granule maturation as shown by cryo-CLEM and polymerase self-interaction shown by SPR. **A)** Examples of cells expressing polymerase-GFP (strain LS5810) and PHB granule stained with Nile Red. i) A tomographic slice from a 3D reconstruction (top left) shows 2 granules in this cell. The polymerase-GFP signal (green) co-localizes with both these granules (bottom left). However, only the larger granule stains for Nile-Red (top right). A composite image shows the co-localization of polymerase-GFP and Nile-Red in yellow in the larger granule (bottom right). PHB granules are outlined in the overlay images by a dotted circle. ii) Another example cell. **B)** Purified polymerase was tested for self-interaction in vitro using surface plasmon resonance. Polymerase was immobilized on the surface using amine crosslinking as ligand, and either full-length polymerase (left graph) or the N-terminal mutant polymerase, which lacks the first 85 amino acids on the N-terminus (right graph), were flown over as analyte at increasing concentrations as indicated.

The condensation of the polymerase to initiate nascent granule formation may be enhanced by polymerase self-interaction. Using surface plasmon resonance, purified PHB polymerase was tested for its ability to self-interact (Fig. 3B left panel). A strong response was observed when increasing concentrations of polymerase are flown over surface-immobilized polymerase, indicating a strong self-interaction of the PHB polymerase.

### The polymerase unstructured N-terminus binds phasin and is required for polymerase expression

We have observed that polymerase condensation precedes the appearance of Nile Red-labeled granules and that 3HB-CoA precursor production, and consequently polymerase catalyzed PHB accumulation, is required for granule formation. We have also observed the continued accumulation of phasin in maturing granules, and an inability of granules to mature in a *phasin* deletion mutant (Fig. 2A and S2A). The inability of granules to increase in size in a *phasin* mutant suggests a defect in maintaining PHB synthesis in the granule. This raises the possibly that there is a promoting effect of phasin on polymerase activity, in turn increasing PHB synthesis. To explain this putative self-reinforcing loop, we studied a potential interaction between polymerase and phasin.

*Caulobacter* polymerase PhaC and phasin PhaP proteins were purified and used in surface plasmon resonance experiments to determine if polymerase and phasin interacted *in vitro*. (SPR, Fig. 4A). With phasin immobilized on the surface, we observed a polymerase concentration-dependent response in signal (Fig. 4A, left graph). This concentration dependent binding was also observed when the polymerase was immobilized on the surface and the phasin flown over (Fig. S5). This direct interaction was surprising, since phasins in two other bacterial species do not directly bind the polymerase when tested by BACTH or crosslink pull-down (39, 40). However, the *Caulobacter* polymerase has an N-terminal, intrinsically disordered domain that is not present in polymerase from either of the two bacteria examined, *P. putida* or *C. necator* polymerase (41). Repeating the SPR with purified, N-terminally truncated polymerase (polymeraseΔN, Table S1) revealed a complete lack of binding with phasin (Fig. 4A, right graph). However, polymeraseΔN was still able to interact with full-length polymerase (Fig. 3B, right graph).

**Figure 4.**
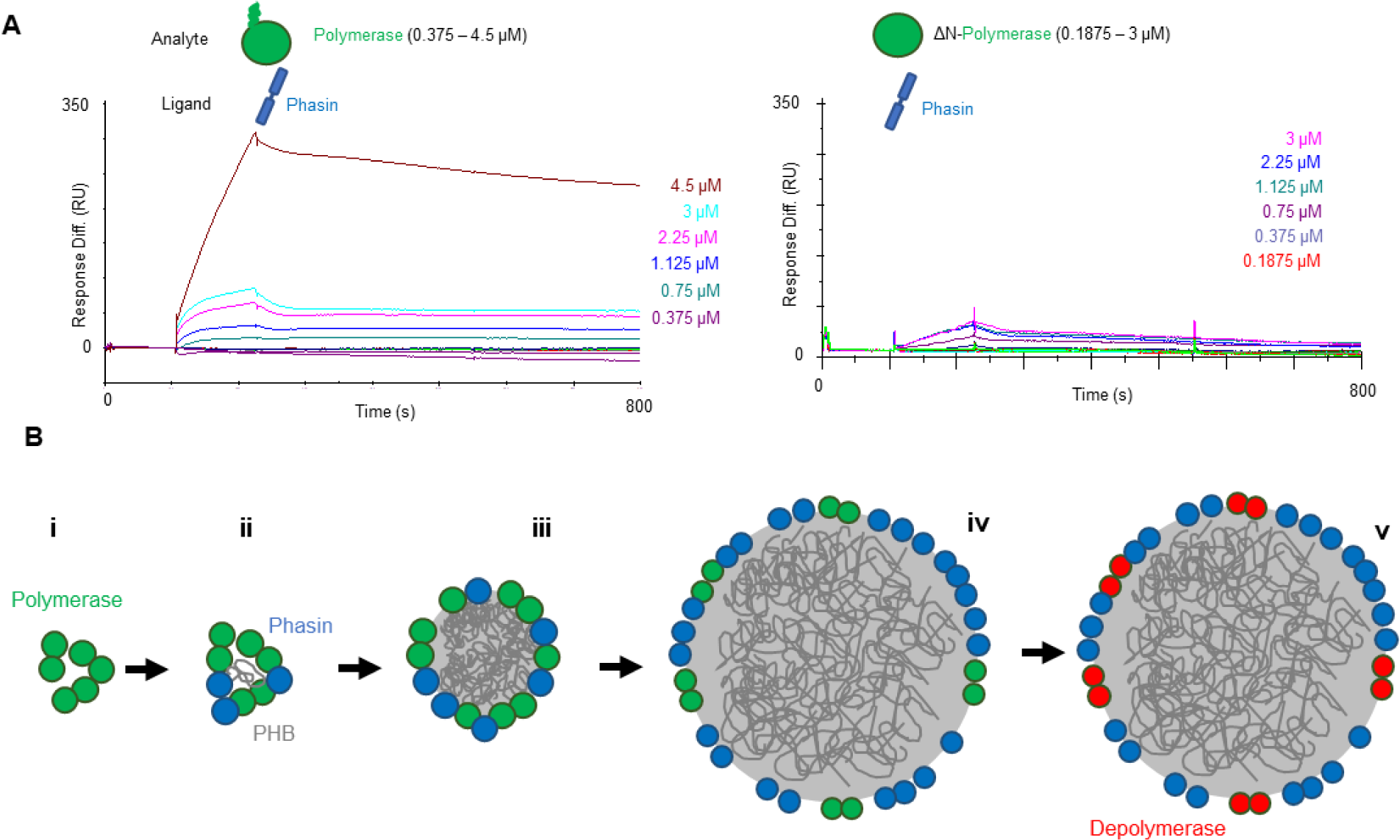
Phasin and Polymerase interact directly through polymerase N-terminus. **A)** Polymerase binds immobilized phasin, but polymeraseΔN does not. Surface plasmon resonance response curves of either polymerase (left graph) or polymeraseΔN (right graph) flown over surface immobilized phasin at increasing concentrations as indicated. **B)** Proposed model for PHB granule formation. i) Polymerase self-interaction (Fig. 3B) promotes condensation that precedes the condensation of phasin (Fig. 2B). ii) Polymerase recruits phasin through direct interaction (Fig. 4A), forming a nascent granule possibly assisted by the early initiation of PHB polymer formation. iii) Ongoing PHB synthesis expands the granule size. iv) As the PHB granule grows until its mature size, more phasin is recruited. v) Once the granule reaches its mature size, the PHB polymerase dissociates from the granule, coincident with the recruitment of the PHB depolymerase (Fig 2C).

## Discussion

In this study, we asked how *Caulobacter* forms PHB granules *de novo*. The PHB polymerase (PhaC) (4), the production of PHB monomers by PhaA and PhaB, and the phasin (PhaP) homologue are essential for the formation and maturation of PHB granules in *Caulobacter* (Fig. 1). Using timelapse fluorescent microscopy and correlative light and cryo-electron microscopy (CLEM), we provide evidence that the composition of PHB granules changes over time as the granule matures (Figs. 2,3). We further used surface plasmon resonance to show that purified PHB polymerase and phasin interact directly through an N-terminal extension of the PHB polymerase that is unique to *Caulobacter.* Based on our results, we propose a model of PHB granule biogenesis and maturation in *Caulobacter* (Fig. 4B): i) Polymerase self-interaction (Fig. 3B) promotes condensation that precedes the condensation of phasin (Fig. 2B). ii) The polymerase can recruit phasin through its N-terminus (Fig. 4A) and form a nascent granule, possibly assisted by the early initiation of PHB polymer formation. iii) Ongoing PHB synthesis expands the granule size and the granule surface. iv) As the granule grows to its mature size, more phasin is recruited to the granule (Fig. 2B). v) Once the granule reaches its mature size, the PHB polymerase dissociates from the granule, coinciding with the recruitment of the PHB depolymerase (Fig 2C).

The sequential localization of granule associated proteins illustrates that the composition of individual PHB granules changes over time, similar to changes in the composition of magnetosomes and carboxysomes during their formation (1, 2, 42). Magnetosomes are membrane bound organelles in which magnetite is stored, enabling their magnetotactic bacterial hosts to orient themselves within a magnetic field, assisting in chemotaxis (1, 43, 44). During magnetosome biogenesis, the bacterial inner membrane invaginates to form a compartment, followed by the accumulation of magnetite (1, 45, 46). Carboxysomes, which carry out carbon fixation and are responsible for efficient photosynthesis under low CO_2_ concentrations, are not encapsulated by a membrane. The key photosynthetic protein, RuBisCo, is first condensed into a pre-carboxysome. Shell proteins are then recruited to encapsulate this condensate generating a finished carboxysome (2, 47). PHB granules are not membrane bound, but there is also no evidence of a continuous protein shell (6, 9). As active PHB synthesis is required for granule formation, PHB ‘seeds’ emerging from the polymerase condensation, possibly aided by condensation of the disordered N terminal portion of the polymerase prior to granule formation, could help drive initial accumulation. Considering PHB is highly water insoluble, phase transition could function as a selective barrier for this organelle. This interface between cytoplasm and PHB is the most likely site for accumulation of granule associated proteins. Since PHB synthesis is required for granule formation, the volume to surface ratio changes as a granule increases in size. This ratio could play a role in selective protein recruitment to the granule surface.

The recruitment of depolymerase (PhaZ) following that of the polymerase answers a previous paradox of futile PHB cycling (48, 49). Published studies observed the expression of both PHB polymerase and depolymerase in cell cultures, suggesting that PHB is both formed and consumed (50, 51). Purified granule populations also show the presence of both polymerase and depolymerase (8). However, in both cases an ensemble measurement was made. In our single cell imaging experiments, we observed that while PHB polymerase and depolymerase can be expressed in the same cell, they do not necessarily co-localize to the same granule at the same time in granule maturation. (Fig 2).

Phasin isolated from *Pseudomonas putida* has been shown to encapsulate PHB *in vitro* (17), and, *in vivo*, phasin protein levels were found to increase during PHB production in *C. necator* and *Caulobacter* (13, 52). *In vitro* reactions of PHB synthesis occur in large (about a centimeter) aggregates indicating the tendency of PHB to self-associate (53). Further, a previous cryo-EM study suggested that the bimodal distribution of granules size might indicate granule fusion *in vivo* (6). Thus, it is possible that PHB granules fuse and that this fusing is coordinated by phasin, although granule fusion has so far not been observed directly. Deletion of *Caulobacter* phasin disrupts the formation of mature granules, and instead leads to the accumulation of what may be small immature granules (Fig. S2). Because of the observed sequential recruitment of polymerase and phasin (Fig 2), the smaller size of PHB granules in a *phasin* mutant could indicate a premature departure of polymerase from this granule. Thus, changes in the number and size of granules likely reflect differences in the rate of granule initiation, granule growth, and departure of polymerase from existing granules. The role of phasin could be to control both the activity and retention of the polymerase at the granule, thereby influencing the final size of mature granules. The direct interaction between phasin and polymerase likely contributes to the retention of polymerase on the granule, in addition to the initial recruitment of phasin by polymerase.

Strong polymerase self-interaction, measured by SPR (Fig 3B), likely contributes to its localized accumulation and the initiation of granule formation. Since PHB synthesis is required for granule formation (Fig 1), cells can control granule formation by directly controlling polymerase activity. Phosphorylation of the polymerase has been reported to control PHB synthesis in *C. necator* (54). However, the reported phosphorylated threonine residue (T373) is not conserved in *Caulobacter.* An alternative method of polymerase regulation could be through degradation. We had limited success detecting a truncated polymeraseΔN version in *Caulobacter* with either Western blot (FLAG-tagged) or fluorescent microscopy (GFP-tagged) (data no shown). However, there was no difficulty purifying these proteins when expressed in *E. coli*. This could indicate the presence of a *Caulobacter* specific degradation pathway to control the polymerase activity, and by extension PHB granule biogenesis.

The polymerase N-terminus directly binds phasin, which may play a role in phasin’s recruitment to the PHB granule, and in retention of the polymerase at the PHB granule by phasin. The intrinsically disordered N-terminal extension of the polymerase is unique to *Caulobacter*, but the N-terminal region shows sequence and functional homology to PhaM in *C. necator*. PhaM contains an intrinsically disordered region comprised of lysine, proline, and alanine repeats (55), which is also found in the *Caulobacter* polymerase N-terminus. It has been reported that *Caulobacter* polymeraseΔN protein lacking this disordered region shows a lag in *in vitro* PHB synthesis (41), like that observed for *C. necator* polymerase (41). Addition of PhaM to the *C. necator* polymerase was reported to overcome this initial lag phase (56), analogous to the absence of lag phase reported for the full length *Caulobacter* polymerase (41). PhaM has been shown to associate with PHB granules, and in mutants lacking PhaM fewer PHB granules are observed (3, 57). Thus, the intrinsically disordered N-terminus of the *Caulobacter* polymerase functions in a manner analogous to the disordered *C. necator* PhaM protein. This underscores a possible role for the N-terminal region of the *Caulobacter* polymerase in the initiation of granule biogenesis. The intrinsically disordered region and polymerase self-interaction may enable localized phase transition which then facilitates the recruitment of the phasin necessary for granule maturation.

PHB granule formation appears to be highly regulated and intricately linked to central carbon metabolism in *Caulobacter*. Precursor synthesis from acetyl-CoA directly links granule formation and growth to the metabolic state of the cell, possibly functioning as an overflow metabolite. If there is no need to make PHB, no organelle will be formed. Phasin is essential for the continued maturation of the PHB granule, potentially by promoting polymerase activity or polymerase retention at the granule. Phasin expression is regulated by the repressor PhaR (13, 58), and PhaR has been reported to bind both DNA and PHB (59). Sequestration of PhaR to the PHB granule has been previously proposed as a mechanism to relieve repression of *phasin* transcription (15), ensuring that phasin is only produced under conditions that promote PHB synthesis. Thus, transcriptional regulation of *phasin* by PhaR ensures proper timing of phasin expression to occur only at the time of granule maturation. In the absence of this additional phasin, the granule is unable to mature, possibly due to premature departure of the polymerase from the granule boundary. As part of the granule boundary, this puts the PHB polymerase at the center of organelle formation; ensuring that its enzymatic activity continues at the granule surface where it directly dictates PHB granule formation and growth, while departure of the polymerase determines the final size of the mature granule.

## Material and Methods

### Bacterial growth conditions

Unless stated otherwise *Caulobacter* strains were inoculated from glycerol stock in PYE for overnight growth at 30°C under aeration. Antibiotics were used for selection during strain construction and added for selective pressure for single-crossover integrated constructs. Antibiotic concentrations that were used (liquid/solid): spectinomycin (25/50 μg/ml), kanamycin (5/25 μg/ml), rifampicin (2.5/5 μg/ml), gentamicin (0.5/5 μg/ml), tetracycline (1/1 μg/ml) and chloramphenicol (2/1 μg/ml). *Escherichia coli* strains were inoculated from glycerol stock in LB for overnight growth at 37°C under aeration. The following concentrations of antibiotics were used for *E. coli*: spectinomycin (50/100 μg/ml), kanamycin (50/50 μg/ml), rifampicin (25/50 μg/ml), gentamicin (15/20 μg/ml), tetracycline (12/12 μg/ml) and chloramphenicol (20/30 μg/ml). All strains used in this study are listed in Table S1.

### Plasmid and strain construction

Plasmids were constructed using previously reported plasmids (60). Plasmids used in this study were either; i) ordered from Synbio Technologies, ii) constructed using Gibson Assembly (GA) protocol from New England Biolabs (NEB) or iii) using restriction-ligation per NEB protocol. Plasmids and primers used in this study are listed in Table S2. Plasmids were isolated using Monarch Plasmid Miniprep kit from NEB and linearized using restriction enzyme by NEB as per manufacturer’s protocol. Inserts were acquired by PCR using KOD Hotstart Mastermix from NovaGen and purified from gel using Monarch DNA Gel Extraction kit from NEB. All cloning was performed in *E. coli* using ultracompetent DH5α from NEB by heat shock as per manufacturer’s protocol. Plasmids were transformed to *Caulobacter* by electroporation. For marker-free integration and deletion, plasmids were first integrated selecting for kanamycin resistance, after which counter-selection on sucrose was performed (60). Kanamycin sensitive candidates were verified by PCR.

pEK001 was constructed by 2 fragment GA using pET28a linearized with BamHI and NheI as backbone and CCNA_01444 amplified from gDNA with EKP001 and EKP002 as insert. pEK003 was constructed by 4 fragment GA using pNTPS138 linearized by HindIII and NheI, *CCNA_02242* CDS without stop codon including 368bp upstream amplified from gDNA genomic DNA (gDNA) with EKP005 and EKP006, mCherry amplified from pCHYC-1 with EKP007 and EKP008, and 800bp downstream of *CCNA_02242* amplified from gDNA with EKP009 and EKP010. pEK005 was constructed by Synbio using PCR cloning, combining synthesized *CCNA_01444* 3’ 800bp into pGFPC-4 backbone. pEK011 was constructed by Synbio using gene synthesis using pET28a as a backbone and *CCNA_02242* as synthesized insert. pEK012 was constructed by Synbio using gene synthesis, combining the fragments 800bp upstream *CCNA_02242*, the chloramphenicol cassette from pMCS-6, and 800bp downstream of *CCNA_02242* into pNPTS138. pEK013 was constructed by Synbio using gene synthesis, combining 800bp upstream of *CCNA_00250*, the tetracycline cassette from pMCS-5, and 800bp downstream of *CCNA_00250* into pNPTS138. pEK014 was constructed by Synbio using gene synthesis, combing 800bp upstream of *CCNA_01444*, the rifampicin cassette from pMCS-3, and 800bp downstream of *CCNA_01444* into pNPTS138. pEK022 was constructed by 2 fragment GA using backbone pXMCS-2 linearized with KpnI and NdeI and insert 5’ 800bp of *CCNA_00544* amplified with EKP015 and EKP016 from gDNA. pEK035 was constructed by amplifying pEK001 using EKP040 and EKP041, digesting the PCR product with NheI and ligating the ends back together. pEK044 was constructed by 4 fragment GA, using pNPTS138 linearized with HindIII and NheI as backbone, 3’ 800bp of CCNA_00250 amplified with MP001F and MP002R from gDNA, mCherry amplified from pCHYC-1 with MP002F and MP002R, and 800bp downstream of CCNA_00250 amplified with MP003F and MP003R from gDNA. pEK052 was constructed by 2 fragment GA, using pTC127 linearized by NdeI and SacI as a backbone, and CCNA_01444 amplified from gDNA with EKP083 and EKP084 as insert. pEK055 was constructed by 2 fragment GA, using pXCHYC-2 linearized with NdeI and NheI as backbone, and CCNA_02242 amplified from gDNA with EKP071 and EKP074 as insert.

### Staining of PHB granules

*Caulobacter* cells were inoculated from glycerol stock into PYE and grown overnight at 30°C. Overnight cultures were subcultured in M2G minimal media and grown overnight at 30°C. For nile red staining, cells were washed in PBS and stained in PBS and 1μM Nile Red (final concentration) for 20 minutes at room temperature. Cells were washed in PBS and subsequently imaged. For Nile Blue staining, 10μM Nile Blue (final concentration) was added directly to the media and cells were stained for 20 minutes at room temperature. Cells were washed in M2G and subsequently imaged.

### Room-temperature fluorescent microscopy and image analysis

Unless otherwise stated, cells were imaged on M2G 1% agarose pads on a Leica Dmi8 multicolor epifluorescence microscope. A phase-contrast 100x oil immersion objective was used with a Lumencor SpectraX LED light source. Images were captured on an EMCCD camera (Hamamatsu C9100 02 CL). GFP was imaged at 480nm excitation and 527nm emission. mCherry, nile red and PHB specific nile blue were imaged at 554nm excitation and 609nm emission. Nile blue membrane stain was imaged at 635nm excitation and 680nm emission. Images were analyzed and quantified using ImageJ (Fiji).

### Cryogenic electron microscopy

Single cryogenic transmission electron micrographs of *C. crescentus* were acquired using a 300 keV electron microscope (G3 Titan Krios^TM^) equipped with direct electron detection camera with a BioQuantum^TM^ Imaging Filter (Gatan, Inc) using a 30-eV slit width. The effective pixel size was ∼6.5 Å, target defocus was 10 µm and the total dose was 25 e^-^/Å^2^. For presentation the data was binned by 4 into 2.6 nm pixels. Acquisitions were controlled using SerialEM software (61).

### Cryogenic correlative light and electron tomography

Cryogenic light microscopy for correlative imaging was acquired on a home-built light microscope (62). Light microscopy was performed using either 488 nm (imaging of GFP channel) or 561 nm (imaging of nile red channel) laser-based illumination through a 100x objective (Nikon CFI TU Plan Apo 100x/NA0.9). Intensities at the sample were kept below 50 W/cm^2^ to avoid sample heating and devitrification(63). Fluorescent emission was detected with an EMCCD camera (Andor iXon) after passing through a 488 nm and 561 nm notch blocking filters and either a 525/50 bandpass for the GFP channel or 607/70 bandpass filter for the nile red channel. Despite both color channels being collected on the same detector, slight shifts in the relative position of the images were corrected for by translating one color channel to align common features, such as grid holes, for each field of view. The excitation beam profiles were measured using a thin film of fluorescent dye embedded in PVA-VA spun on a coverslip. Fluorescent brightness across the field of view was corrected for by dividing acquired images by a Gaussian fit to the measured beam profiles.

Subsequent electron tomography was performed on a 300 keV electron microscope (G3 Titan Krios^TM^) equipped with direct electron detection camera with a BioQuantum^TM^ Imaging Filter (Gatan, Inc) using a 20-eV slit width. The effective pixel size was 4.49 Ångstroms. The tilt series was acquired unidirectionally over ±46 degrees in 2 degree steps. Total dose was 100 e^-^/Å^2^. Reconstructions were performed using IMOD with data binned-by-4 to produce 1.8 nm effective pixels (64). Registration of light microscopy and electron tomography datasets was performed using a projective transformation first to a lower magnification cryoEM micrograph and then a similar transformation to the higher magnification tomography datasets. Further details on the registration process can be found in Dahlberg et al (38).

### Timelapse microscopy

Overnight PYE cultures of *Caulobacter* were subcultured in M2G minimal media and grown at 30°C overnight. For xylose induction, the cells were pelleted and resuspended in M2G+0.1% xylose and spotted on M2G+0.1% xylose pads. For glucose recovery, the culture was washed in M2 and resuspended in M2 and starved 24 hours at 30°C. Just prior to imaging, cells were pelleted and resuspended in M2G and spotted on M2G pads. The coverslip was sealed by wax to prevent evaporation and the recovering cells were imaged at 10-minute (xylose induction) or 15-minute (starvation recovery) intervals for at least 3 hours.

### Protein purification

His-tagged proteins were expressed in *E. coli* BL21(DE3) using the pET28a system (polymerase = pEK001, phasin = pEK011 and polymeraseΔN = pEK035). Cells were grown overnight at 37°C in LB and diluted 1:400 into prewarmed LB. At an OD_600_ of approximately 0.5, protein expression was induced with 0.4μM IPTG and the culture was shifted to 18°C for another 3 hours. Cells were washed in ice-cold PBS and the pellet was stored at -80°C. Cells were thawed on ice and resuspended in ice-cold lysis buffer (50mM HEPES-KOH pH 8.0, 500mM KCl, 25mM imidazole, 1 protease tablet per 50mL buffer, 200U benzonase nuclease at 0.3ul per 1L culture). Cells were lysed by sonication (3 sec pulse, 12 sec rest, 100% amplitude, 4 minute total pulse) and the lysate was cleared by centrifugation (30 minutes, 4°C at 29000 rcf). Cleared lysate was inoculated with pre-equilibrated Ni-NTA resin (1ml per 1l culture) for 2 hours at 4°C under gentle agitation. The lysate-resin mixture was separated by a gravity column (Poly-Prep Chromatography Column, Bio-Rad) and the resin was washed with 15 column volumes (CV) wash buffer (50mM HEPES-KOH pH 8.0, 500mM KCl, 25mM imidazole). The sample was eluted in 5 CV elution buffer (50mM HEPES-KOH pH 8.0, 500mM KCl, 250mM imidazole, 10% glycerol) and stored at -80°C. For the removal of the His6-tag, a Thrombin CleanCleave Kit (Sigma) was used.

### Surface Plasmon Resonance

Reagents: Sensor CM5 chips, N-Hydroxysuccinimide (NHS), 1-Ethyl-3-(3-dimethylaminopropyl) carbodiimide (EDC), and ethanolamine hydrochloride (EA) were obtained from GE Healthcare/Cytiva. Surface Plasmon Resonance experiments were performed on a BIACORE 3000 biosensor system (GE Healthcare/Cytiva) at 25°C. Proteins were dialyzed against Assay Buffer (20mM HEPES pH7.4, 200mM NaCl) before use. The different proteins (His-tagged or untagged Polymerase (PhaC) and phasin (PhaP)) were covalently immobilized on the surface of a CM5 biosensor chip (GE Healthcare/Cytiva) by amine coupling chemistry using N-hydroxysuccinimide (NHS) and N′-(3-dimethylaminopropyl) carbodiimide hydrochloride (EDC) according to the manufacturer’s instructions. To investigate binding of Polymerase (PhaC) or phasin (PhaP) to the immobilized Polymerase (PhaC) or phasin (PhaP), dialyzed proteins were diluted in Assay Buffer and injected over the Ligand containing surface for 2min at different concentrations at a flow rate of 30 μl/min. For each experiment at least 5 different concentrations of analyte proteins were injected over each experimental and control flow cell. Dissociation was allowed to occur at the same flow rate for 300 sec. followed by Regeneration Buffer (20mM HEPES pH7.4, 1M NaCl) at a flow rate of 100 μl/min for 30sec and Assay Buffer alone at a flow rate of 30 μl/min to allow the baseline to stabilize. All data were corrected for unspecific binding by subtracting the signal measured in a control cell lacking immobilized ligand. Data analysis was performed using the BIAevaluation software 4.1 (GE Healthcare/Cytiva).

## Acknowledgements

We thank T. Chong for providing plasmids and thank T. Chong and S. Saurabh for helpful discussions on experimental aspects of the manuscript. We thank all members of the Shapiro laboratory for constructive feedback throughout this work.

## Author contributions

This study was conceived by E.A.K. Experiments were designed by E.A.K., with input from L.S., M.E. and P.D.D. Strains for this study were engineered by E.A.K. and M.P. Room temperature fluorescent microscopy was performed by E.A.K. and M.P and analyzed by E.A.K. Protein purification and BACTH were done by E.A.K. SPR was performed by J.T. and M.R.E. and analyzed by M.R.E. and E.A.K. Cryo-EM and CLEM measurements were performed and analyzed by P.D.D. and E.A.K. The manuscript was written by E.A.K. and L.S., with input from P.D.D. and M.R.E.

## Data and material availability

Plasmids and strains used and created for this work can be obtained on request.

---

**Figure S1.**
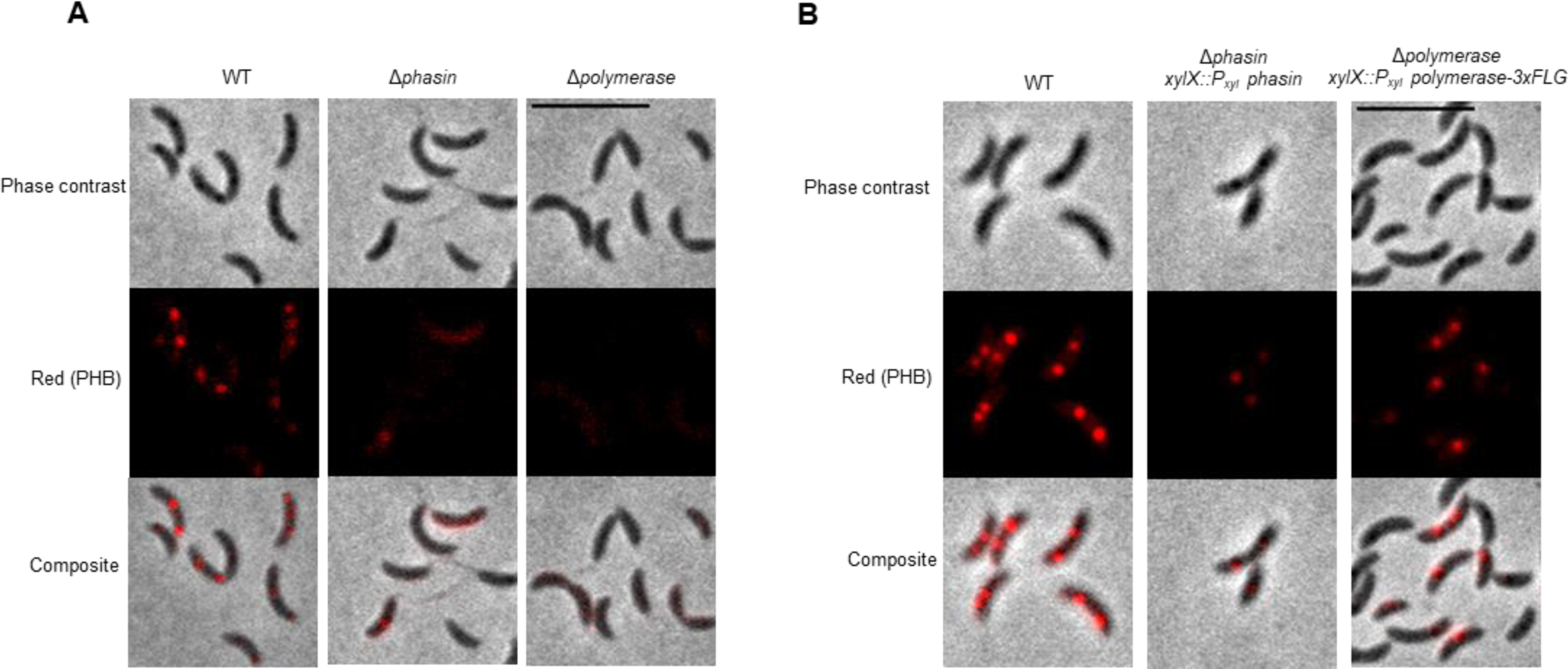
WT and mutants stained by nile blue dye. **A)** Deletion of either polymerase PhaC or phasin PhaP results in a loss of nile-blue stained granules. Strains used are WT, LS5812 and LS5813. Cells are incubated for 20 minutes with Nile Blue prior to imaging. Nile Blue stained PHB granules can be visualized in the red channel (554nm/609nm excitation/emission). Scale bar is 5 μm. **B)** Complementation of polymerase and phasin deletion mutants. A copy of either *phasin* or *polymerase-3xFLAG* was expressed from the xylose locus (*xylX*) using a xylose inducible promoter. Cells were grown in presence of 0.1% xylose in M2G media and PHB granules were visualized using Nile-Blue staining. Strains used are WT, LS5826 and LS5872. Scale bar is 5 μm.

**Figure S2.**
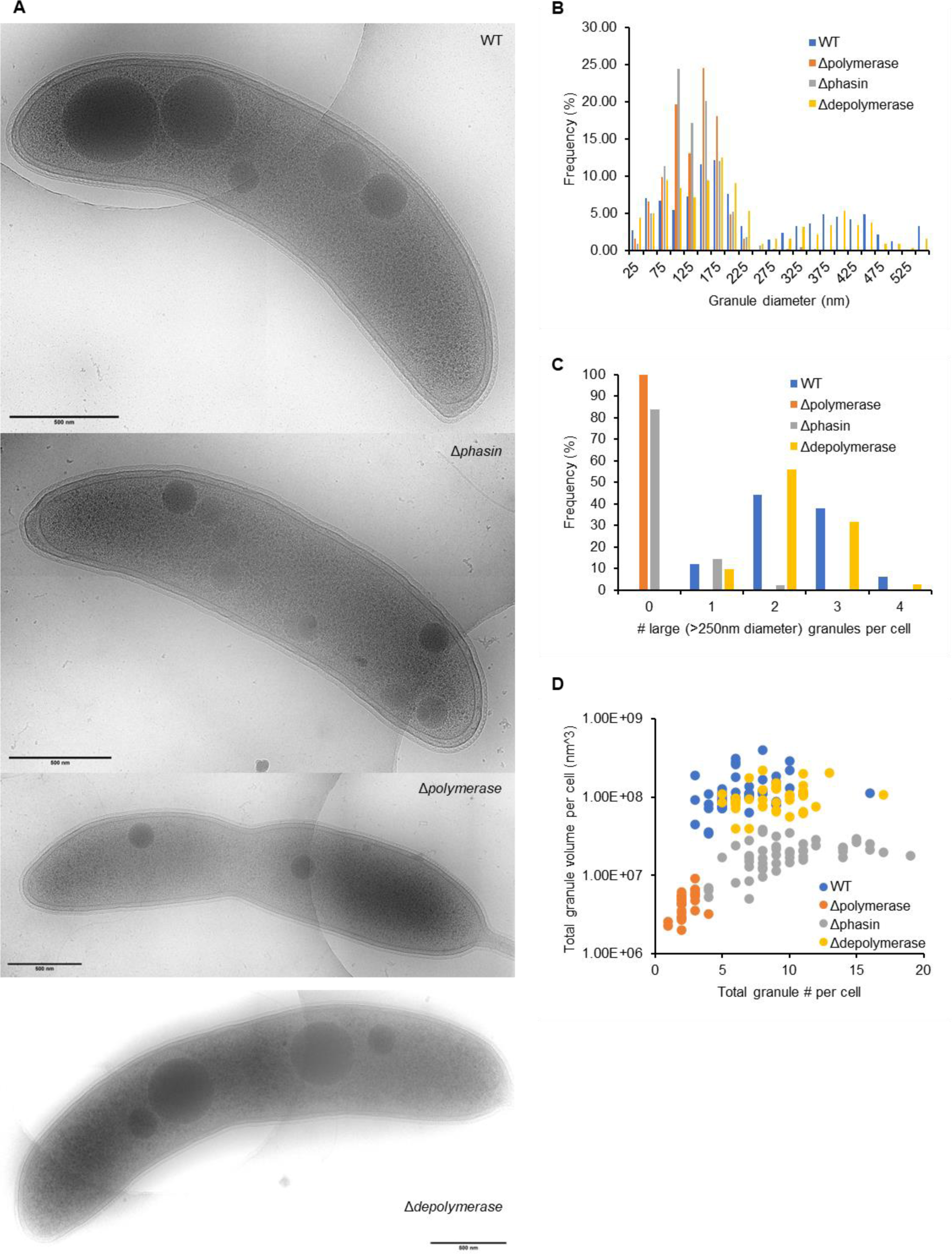
Granule quantification by cryo electron microscopy. **A)** Cryo electron micrographs of WT (top), Δ*phasin* (middle-top, LS5812), Δ*polymerase* (middle-bottom, LS5813) and Δ*depolymerase* (bottom, LS5811) cells. Scale bar is 500 nm. **B)** Frequency of granule diameters measured from 25-50 cells per sample. All granules are measured, so this includes both PHB and poly-P granules. Only poly-P granules were observed in the Δ*polymerase* cell. **C)** Frequency of granule diameters for granules with a diameter >250nm. **D)** Total cumulative granule volume per cell. Granule volumes were calculated based on granule diameters, and all granule volumes were summed per cell. Although Δ*phasin* cells have more granules per cell than WT (9.8 vs 6.6 granules per cell), the total granule volume per cell is 6.6x lower than WT. In EM images, both PHB and poly-phosphate granules can be found in WT cells. Although poly-phosphate granules have been reported to be more electron dense than PHB granules (6), we were unable to quantify such a difference in our images. Since cells lacking the PHB polymerase will only contain poly-phosphate granules, we used this as the expected average number of poly-phosphate granules per cell. In Δpolymerase cells there were on average 2.3 granules per cell at an average 143nm diameter. We observed no granules with a diameter over 250nm in Δpolymerase cells, which we assume will be the maximum diameter for poly-phosphate granules. WT cells contain both PHB and poly-P granules, on average 6.6 granules per cell. On average 2.4 granules per cell have a diameter that exceeds 250nm, comparable to the average of 2.1 Nile Red stained granules per cell (Fig. 1D), which should only stain PHB granules. The remaining granules (on average 4.2 per cell) that do not exceed 250nm therefore are likely a mix of poly-phosphate granules and smaller PHB granules. For the phasin mutant we observe on average 9.8 granules per cell, of which only 0.2 granules per cell exceed 250nm diameter. This subset (16.3%) of the cells reflects the 7.3% cells containing Nile red stained granules (Fig. 1D).

**Figure S3.**
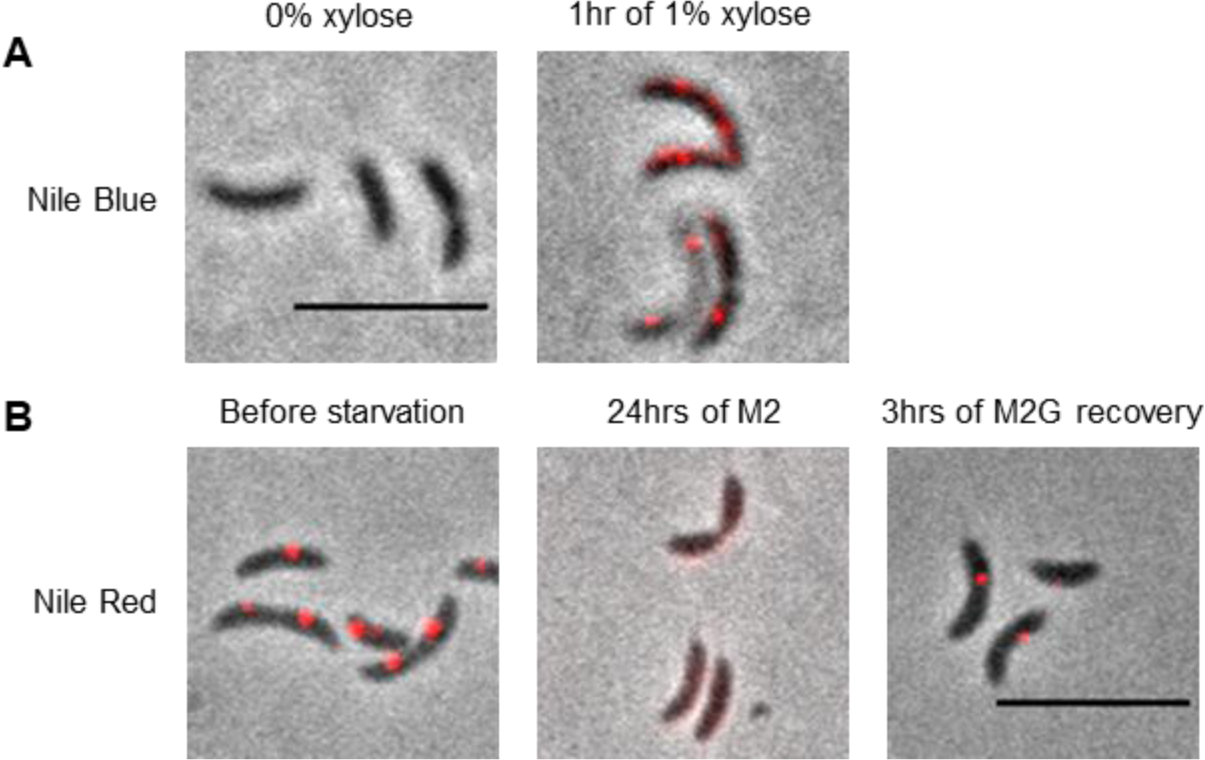
Controlled granule formation in *Caulobacter*. **A)** Cells with PhaAB expression under xylose control (LS5814) do not show any Nile Blue puncta in the absence of xylose (left). Addition of 1% xylose results in clear Nile Blue puncta after only 1 hour of growth (right). **B)** WT cells grown into stationary phase in M2Glucose show clear Nile Red puncta (left). Replacing the media with M2 and starving the cells for 24 hours results in a loss of Nile Red puncta (middle). Replacing the media again with M2Glucose recovers the Nile Red puncta after 3 hours (right). Scale bars are 5μm.

**Figure S4.**
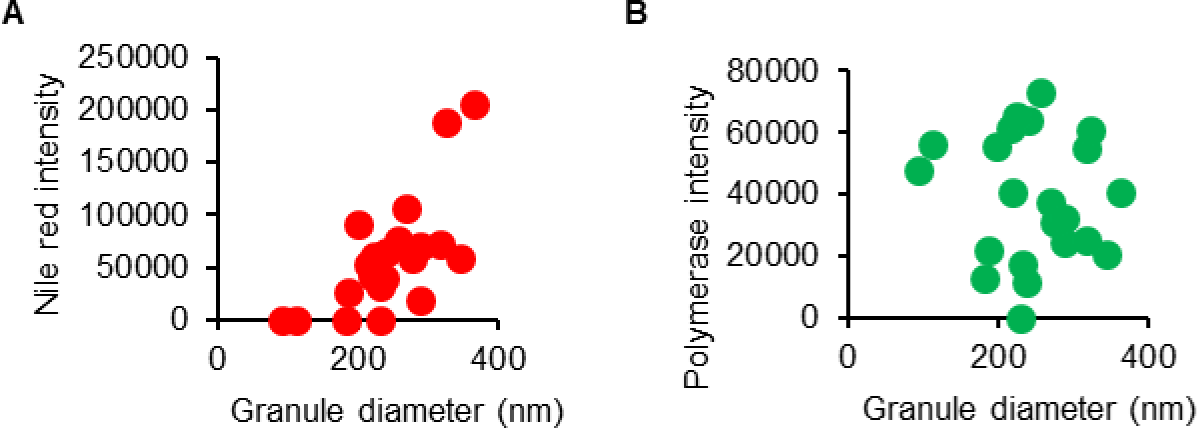
Nile Red intensity correlates to PHB granule size. Quantification of **A)** Nile Red and **B)** Polymerase-GFP signal of cryo-CLEM images. Fluorescent intensities were measured for 22 granules and compared to the diameter of these granules. Only granules that had Nile Red and/or GFP signal were analyzed. A positive correlation is observed between granule diameter and Nile Red intensity (A), while no correlation is seen between granule diameter and polymerase-GFP signal.

**Figure S5.**
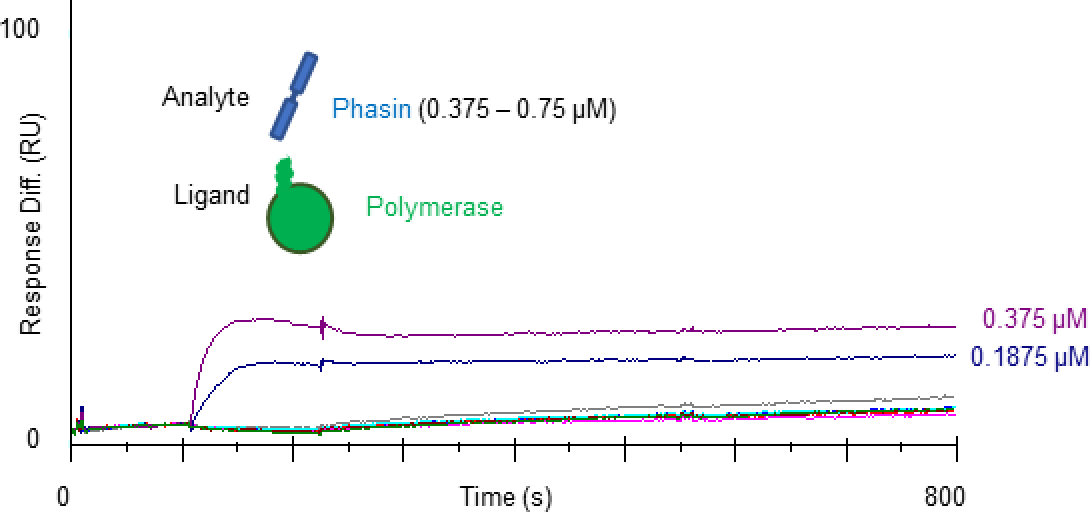
Phasin binds immobilized Polymerase in SPR. Surface plasmon resonance response curves of increasing concentrations of phasin flown over surface immobilized polymerase at increasing concentrations as indicated.

**Table S1.** Bacterial strains.

A. List of *Caulobacter* strains used in this study. Plasmids integrate either by single- (:) or double- (::) crossover.
| Name | Genotype | Source |
| --- | --- | --- |
| LS101 (WT) | NA1000/CB15N | Lab strain |
| LS5811 | CCNA_00250::tetR | LS101::pEK012 |
| LS5813 | CCNA_01444::rifR | LS101::pEK014 |
| LS5812 | CCNA_02242::catR | LS101::pEK013 |
| LS5814 | CCNA_00544:P<sub>xyl</sub>-CCNA_00544 kanR | LS101:pEK022 |
| LS5810 | CCNA_01444:CCNA_01444-GFP gentR | LS101:pEK005 |
| LS5809 | CCNA_02242::CCNA_02242-mCherry | LS101::pEK003 |
| LS5816 | CCNA_02242::CCNA_02242-mCherry, CCNA_01444:CCNA_01444-GFP gentR | LS5809:pEK003 |
| LS5871 | CCNA_02242::CCNA_02242-mCherry, CCNA_01444:CCNA_01444-GFP gentR, CCNA_00544:P<sub>xyl</sub>-CCNA_00544 kanR | LS5816::pEK022 |
| LS5819 | CCNA_00250::CCNA_00250-mCherry | LS101::pEK044 |
| LS5821 | CCNA_00250::CCNA_00250-mCherry, CCNA_01444:CCNA_01444-GFP gentR | LS5819:pEK003 |
| LS5872 | CCNA_02242::catR, xylX:P<sub>xyl</sub> CCNA_02242 kanR | LS5812:pEK055 |
| LS5826 | CCNA_01444::rifR, xylX:P<sub>xyl</sub>-CCNA_01444-3xFLAG kanR | LS5813::pEK052 |

B. List of *E. coli* strains used in this study.
| Name | Source |
| --- | --- |
| Ultracompetent DH5α | New England Biolabs |
| BL21(DE3) | Novagen |

**Table S2.**
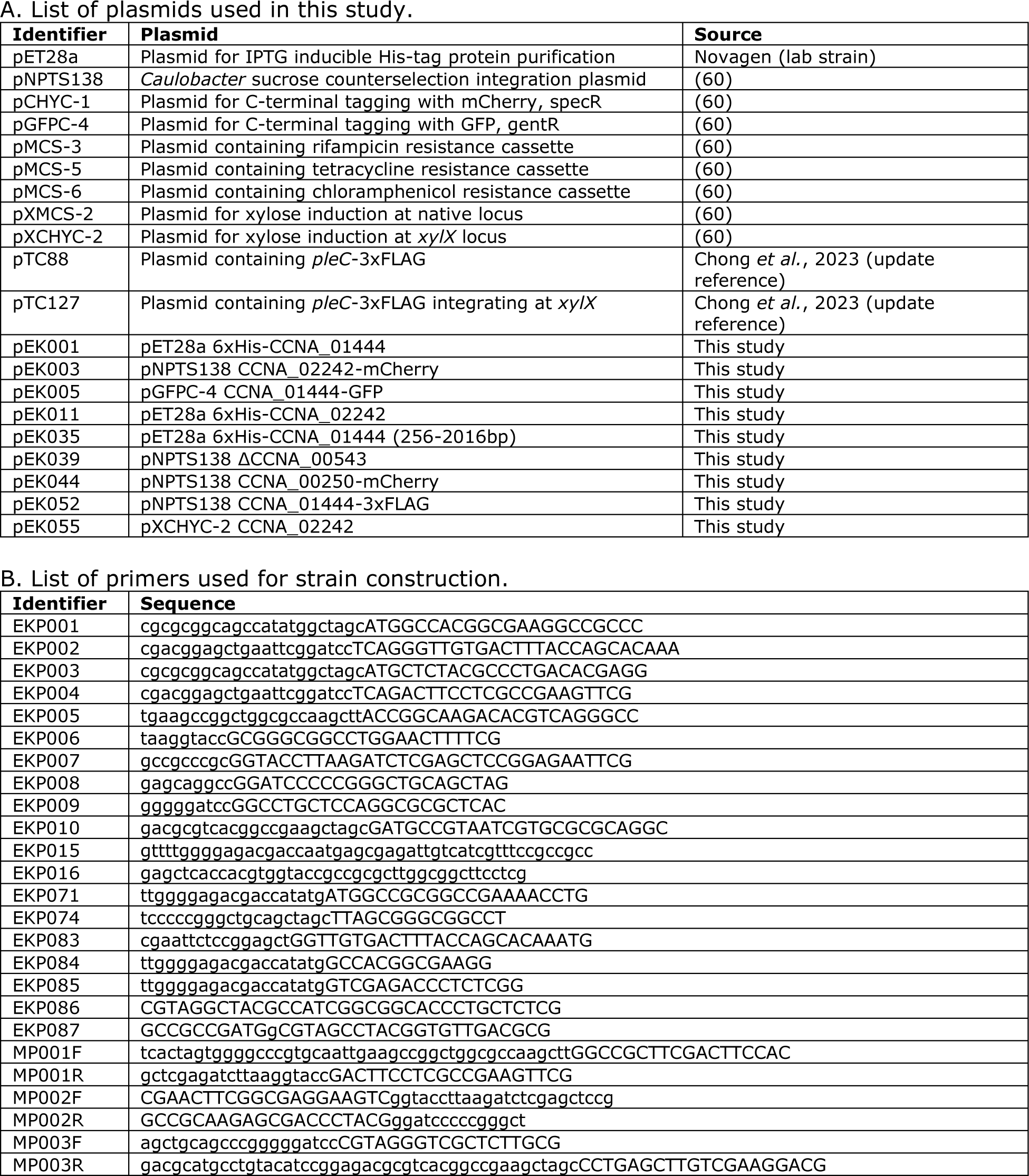
Plasmids and primers.

